# Compositional decoding of neural activity enhances generalization in handwriting BCIs

**DOI:** 10.64898/2026.02.09.704932

**Authors:** Sowmya Manojna Narasimha, Jingya Huang, Ram Dyuthi Sristi, Vikash Gilja, Gal Mishne

## Abstract

Recent brain-computer interfaces (BCIs) have achieved state-of-the-art performance in decoding behavior from neural activity. These models are typically trained on a constrained set of behaviors, which limits their ability to generalize to real-world settings where behavior is variable, complex, and context-dependent. However, many complex behaviors can be decomposed into a set of reusable behavioral motifs, indicating a compositional organization. Here, we analyze human intracortical neural activity underlying attempted handwriting and find signatures of neural compositionality at a finer resolution than individual letters. We further introduce a compositional temporal decoding model, MOtif-based Temporal Inference Framework (MOTIF), that jointly predicts the fine-scale behavioral motifs (e.g., strokes, phonemes) and the longer-timescale behavior class (e.g., characters, words). We show that the compositional structure leveraged by MOTIF enables improved generalization in few-shot learning. Our results demonstrate that explicitly incorporating compositionality into neural decoders can enhance generalization and sample efficiency, while providing a principled approach to designing more scalable, robust, and interpretable BCIs.

## I. Introduction

Brain-computer interfaces (BCIs) aim to restore communication and motor function for individuals with paralysis by decoding intended motor behavior from neural activity. Advances in intracortical recording technologies and neural decoding algorithms have led to substantial improvements in BCI performance. In particular, recent works in speech and handwriting BCIs have achieved state-of-the-art decoding accuracies, with communication rates approaching those of natural human communication [1]–[5]. These advances highlight the potential for BCIs and neural decoders to serve as assistive technologies suitable for daily use.

Despite this progress, most existing BCIs are trained in controlled experimental settings on a small set of constrained and repetitive behaviors. While this paradigm facilitates model training and benchmarking, it limits their ability to generalize to naturalistic settings. In real-world settings, behavior is variable, context-dependent, and structurally complex [6]. Yet, complex natural behavior reflects compositional organization, i.e. it can be decomposed into a set of reusable motifs [7]–[11]. For example, in speech, sequential combinations of phonemes form words, and in handwriting, strokes form characters. Recent brain-to-text BCIs for attempted speech have leveraged compositionality by decoding phoneme sequences to reconstruct words, achieving record communication rates [2]–[4].

However, outside of speech, compositional modeling for human BCI has been relatively unexplored, and it remains unclear whether similar principles apply to other motor behaviors that engage non-speech motor cortical regions [1], [10], [12]. In this work, we examined whether the neural activity underlying handwriting [1] exhibits compositional structure and whether explicitly modeling this structure into decoders improves generalization. We first analyzed population neural activity during attempted handwriting to assess the presence of shared, reusable neural motifs across multiple characters. We found signatures of motif compositionality in neural activity, with distinct temporal segments across characters reusing similar neural patterns.

To leverage the observed compositionality in neural activity, we proposed a compositional temporal decoding model, MOtif-based Temporal Inference Framework (MOTIF), that jointly predicts motifs (strokes or phonemes) and behavior classes (characters or words). We benchmarked MOTIF against baseline models that perform behavior classification alone. On a 50-50 test-train split, both models achieved comparable performance, with MOTIF additionally identifying the compositional motifs. In a two-shot generalization test, MOTIF significantly outperformed the baseline model in the held-out class. Our results suggest that compositional neural decoders improve on sample efficiency, generalizability, and robustness. By decoding behavioral motifs from neural activity, this work outlines a principled strategy for designing BCIs that generalize to naturalistic settings.

The rest of the paper is organized as follows: Section II describes the neural data used in the study, our proposed model (MOTIF), and evaluation metrics. Section III presents our analysis of compositional structure in handwriting neural activity, comparison between MOTIF and baseline model performance, and interpretability analysis of MOTIF. Finally, Section IV summarizes the work and discusses the implications of these findings for the design of generalizable and interpretable BCI decoders.

## II. Methods

### A. Neural Data

We evaluated model performance on publicly available intracortical neural recordings from human participants performing attempted handwriting of single characters [1] and attempted speech of words [2].

Handwriting neural activity [1] is obtained from participant T5, who has a high-level spinal injury and is paralyzed neck-down. Two Utah microelectrode arrays with a total of 192 electrodes are implanted in the hand motor region. In this task, T5 is visually prompted to attempt writing single lowercase English letters per trial, with an instructed delay between the visual cue and the movement go-cue. Because of T5’s paralysis, ground-truth kinematic measurements (e.g., hand-position trajectories) are unavailable, and only the identity of the prompted letter is known for each trial. We analyzed neural activity obtained across multiple trials on a single day. The dataset comprised 27 repetitions of each of the 26 letters in the English alphabet. Threshold-crossing neural activity is Gaussian-smoothed (50ms, 5 timepoints) and time-aligned to the go-cue across all trials prior to decoding.

Speech neural activity [2] is obtained from participant T12, who has ALS, and is limited in her ability to vocalize and produce orofacial movements. Neural activity is recorded using two microelectrode arrays (128 channels in total) implanted in the left Ventral Premotor Cortex. As with the handwriting dataset, precise articulatory behavioral measurements are unavailable, with only the prompted word identity known for each trial. In addition to threshold-crossing neural activity, spike band power is also included for decoding, yielding a 256-dimensional neural feature vector at each timepoint. We analyzed neural activity obtained across multiple trials in a single day, with 20 repetitions per word in the 50-word dataset [13]. Neural activity is Gaussian-smoothed (80ms, 4 timepoints) and time-aligned to the go-cue across all trials prior to decoding.

### B. Compositional Motifs Identification

We defined the compositional strokes that constitute a letter and determined the phonemes that constitute a word. These strokes and phonemes serve as the compositional motif classes used for motif classification in our proposed decoding model. These motifs are based solely on the behavior studied (handwriting, speech) and are determined independently of the underlying neural activity.

Handwriting across individuals exhibits substantial variability in the selection and ordering of strokes used to produce the same final characters. Since ground-truth kinematic trajectories are unavailable in the attempted handwriting data [1], we adopt a canonical stroke representation for each character. Stroke motifs were defined using kinematic templates provided in [1], which followed T5’s verbal description of how he would write the character (Fig. 1). Analysis of the templates for the 26 letters in the English alphabet yielded 10 distinct strokes, whose combinations were sufficient to represent all letters.

**Fig. 1.**
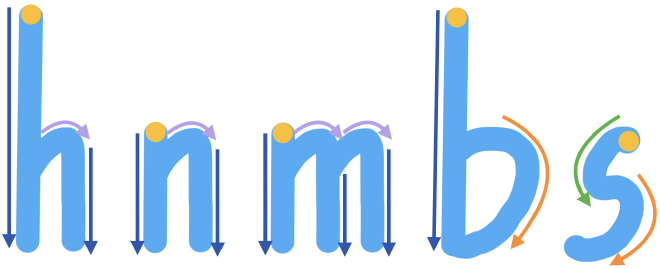
Stroke Composition of Letters. Sample stroke composition of the letters ‘h’, ‘n’, ‘m’, ‘b’, and ‘s’ based on handwriting templates [1]. Distinct stroke motifs are indicated by different colors. The yellow dot represents the start of the handwriting trajectories.

The compositional phonemes for a given word were determined using the CMU Pronunciation Dictionary [14], which provided the standard North American pronunciation for words. All 50 words could be expressed as a combination of 33 distinct phonemes.

### C. MOtif-based Temporal Inference Framework (MOTIF)

Our proposed compositional model, MOTIF, is built on a Temporal Convolutional Network (TCN) [15], [16] backbone, and comprises a motif-prediction branch and a behaviorprediction branch (Fig. 2).

**Fig. 2.**
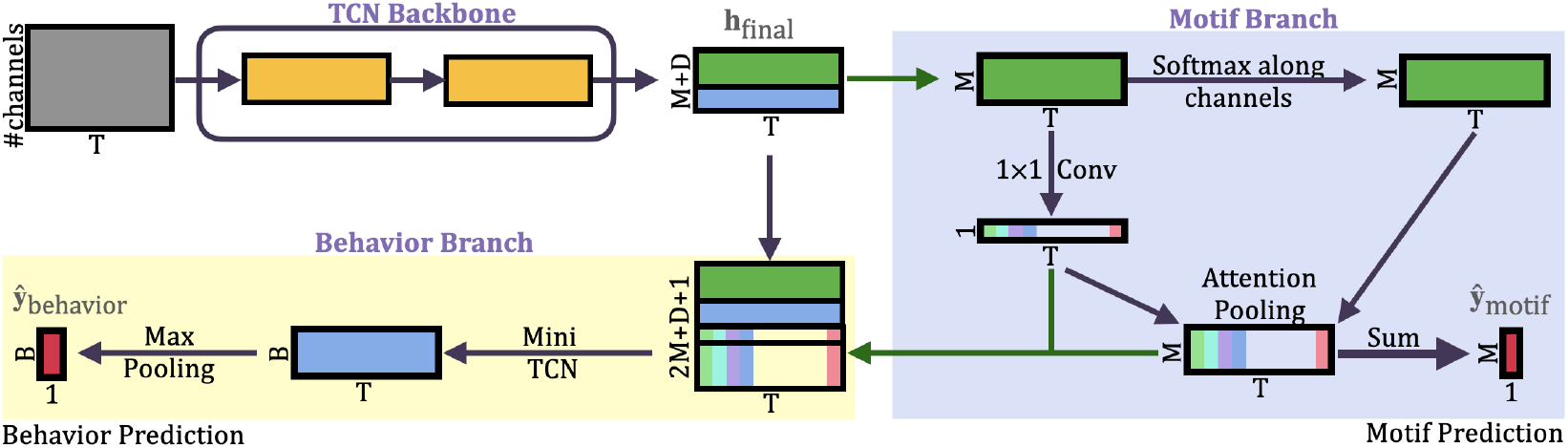
**MOtif-based Temporal Inference Framework (MOTIF)** comprises a TCN backbone, a motif prediction branch (on the right, background highlighted in blue), and a behavior prediction branch that integrates motif representations (on the left-bottom, background highlighted in yellow).

The TCN backbone is used to capture temporal structure in the neural activity. The backbone consists of a stack of causal temporal convolution (TC) blocks, each comprising two temporal convolutional layers with a residual connection from the first layer to the second. The TC layer performs the convolution operation only along the time dimension, and the filters identify temporal structure in neural activity [17], [18]. The output 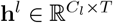 of the *l*-the layer with *C*_*l*_ channels is given by

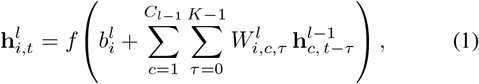

where *K* is the kernel size, 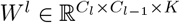 are the kernels, 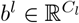 the bias and *f* (·) represents non-linearity.

Within each block, the second convolutional layer uses a dilation factor of 2, enabling the network to capture temporal dependencies over longer durations. Causal convolutions ensure that the latent representation at each timepoint depends only on current and past inputs, preventing information leakage from future data. The final backbone output, denoted **h**_final_, is an input to both the motif prediction and behavior prediction branches.

The *motif prediction branch* identifies the short-timescale behavioral motifs (e.g., strokes or phonemes) present in a trial. The first *M* channels of the TCN backbone output (**h**_final,1:*M*_ ∈ ℝ^*M ×T*^) are passed as input to this branch, where *M* is the number of motifs classes present in the dataset and *T* is the number of timepoints. Softmax is applied along the channel dimension at each timepoint, encouraging the model to select a single dominant motif class per timepoint. A 1 *×* 1 convolution is applied on **h**_final,1:*M*_, producing *attention* weights that indicate the contribution of neural representation at each timepoint towards motif prediction. Attention pooling is then applied along the temporal axis by weighting the softmaxed features at each timepoint with their corresponding attention weights. This enables the model to predict the motifs by selectively attending to specific timepoints. The attention-pooled output is summed across the temporal axis to produce motif logits. Motif predictions (**ŷ**_motif_ ∈ {0, 1} ^*M*^) are determined through independent binary classification for each of the *M* motifs.

The *behavior prediction branch* predicts longer-timescale behaviors (e.g., character or word) in each trial. This branch leverages the TCN backbone output (**h**_final_), the attention weights, and the attention-pooled motif features, thereby informing behavioral prediction with motif-level structure. As the representations learned by the motif branch do not differentiate between behaviors with identical compositional motif sets (e.g., ‘h’ and ‘n’), the final *D* channels in **h**_final_, that are not passed to the motif prediction branch, facilitate distinguishing such behaviors. The concatenated features are then passed through a causal temporal convolutional layer with *B* output channels, where *B* is the number of behavioral classes present in the dataset. Finally, max pooling across the temporal axis is applied to obtain the behavior prediction (**ŷ**_behavior_).

MOTIF is trained using a multi-objective loss that jointly optimizes motif- and behavioral-level predictions. Independent binary cross entropy loss is calculated for each motif class, and the average loss across motifs is considered as the motif loss ℒ_motif_. The behavioral loss is optimized using categorial cross entropy ℒ_behavior_. The overall MOTIF training objective combines the two losses: ℒ_compositional_ = *λ*ℒ_motif_ + ℒ_behavior_, where the hyperparameter *λ* balances fine-resolution motif loss and longer-timescale behavioral loss. In contrast, the behavior-only prediction model is trained only on the behavioral loss, ℒ_baseline_ = ℒ_behavior_.

### D. Evaluation

The models are evaluated on two settings: a 50-50 test-train split and a 2-shot generalization test. The 50-50 test-train split is used to assess performance when sufficient training data is available to the models. Models that perform well under standard generalization may still suffer from insufficient sample representation, particularly when inputs such as neural activity are intrinsically variable [18]–[22]. To further probe robustness in low-data and noisy neural regimes, we evaluate model generalization using a two-shot generalization test. In this setting, only two trials of a target behavior class are included in the training data, while a 50-50 test-train split is maintained for all the remaining classes.

The metrics used to evaluate the model performance are behavioral prediction accuracy, partial motif accuracy, exact motif accuracy, and motif Jaccard score. Partial motif accuracy measures the mean motif-level agreement between the model prediction and the ground truth, and is defined as follows:

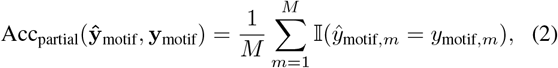

where 𝟙(·) is the indicator function. Exact motif accuracy is a stricter metric that requires exact prediction of the complete motif set and is defined as follows:

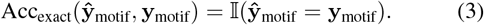

Motif Jaccard score is computed as the size of the intersection over the union of the predicted and ground-truth motif sets and is defined as follows:

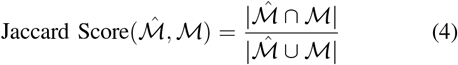

where, 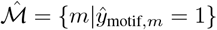 and ℳ = {*m*|*y*_motif,*m*_ = 1}.

### E. Implementation Details

All models were implemented in PyTorch [23] and trained using the Adam optimizer [24]. Model hyperparameters, including the number of layers in the TCN backbone, kernel sizes, channel dimensions, and *λ* for the motif and behavioral losses, were selected based on validation performance. These hyperparameters were maintained across MOTIF and baseline models to ensure fair comparison. All models were trained using CUDA-accelerated implementations on NVIDIA Quadro RTX8000 GPUs. The reported results are the average model performance across five random initializations.

## III. Results

### A. Evidence of Compositionality in Handwriting Neural Data

We first examined whether neural activity underlying handwriting exhibited signatures of compositionality. We analyzed population neural activity to detect shared, reusable neural motifs across multiple behavioral classes. Because the same behavioral motif could occur at different temporal positions across behaviors, timepoints across behaviors that were close in neural space were identified. For each character *i*, at timepoint *t*, the similarity between its trial-averaged neural state 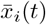 and that of all other characters 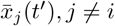 across all timepoints *t*^*′*^ ∈ [0, *T*] is calculated. At each timepoint *t*, only the 100 nearest neighbors 𝒩_*i*_(*t*) of behavior *i* in the population activity space are considered, yielding a sparse spatio-temporal similarity matrix *S*_*i,j*_ ∈ ℝ^*T ×T*^ :

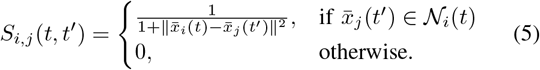

Spatio-temporal neural similarity matrices were computed between the neural activity underlying all pairs of letters. The nonzero values in *S*_*i,j*_ correspond to timepoint pairs with highly similar neural patterns across characters, thereby identifying segments of activity that are reused across behaviors. Analysis of the similarity matrices revealed structured bands of elevated neural similarity (Fig. 3). These bands corresponded to temporally localized neural activity patterns that recur across letters sharing common stroke components. The temporal positioning of these bands varied across letters, reflecting differences in the ordering and duration of the underlying strokes.

**Fig. 3.**
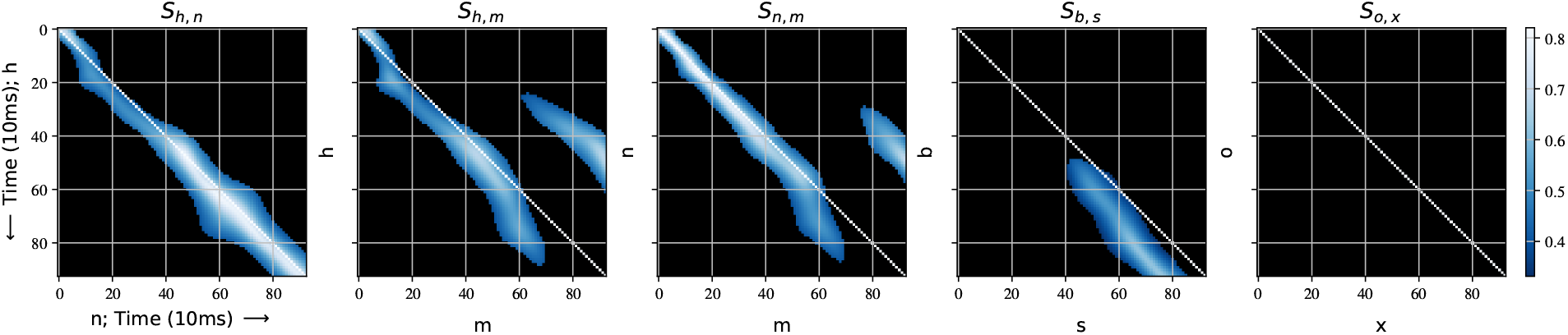
Neural Similarity Matrices. Similarity matrices between neural trajectories corresponding to the characters (h, n), (h, m), (n, m), (b, s), and (o, x). Distinct neural similarity banding is seen in the first four similarity matrices. Double banding is observed in *S*_*h,m*_ and *S*_*n,m*_. A similarity band in the lower right quadrant is observed in *S*_*b,s*_. No similarity bands are present in *S*_*o,x*_. The main diagonal is highlighted for reference.

Representative examples illustrate this structure. For instance, the neural similarity matrix *S*_*h,n*_ exhibits high similarity bands along the main diagonal, consistent with the shared stroke components: vertical downward (↓) and hook (↷) strokes. Slight deviations of similarity bands along the main diagonal in the earlier timepoints corresponding to *h* reflect differences in the temporal extent of the shared strokes, i.e., longer vertical downward stroke in *h* when compared to *n*. In contrast, the similarity matrices *S*_*h,m*_ and *S*_*n,m*_ exhibit multiple parallel bands, indicating the reuse of the neural activity potentially associated with the hook (↷) stroke. As observed earlier, slight deviations along the main diagonal are consistent with the temporal extent of shared strokes.

Other letter pairs exhibit more selective similarity. For example, *S*_*b,s*_ only contains a similarity band in the lower-right quadrant, corresponding to the shared clockwise arc (⟳) stroke occurring in the latter half of the characters *b* and *s*. The absence of similarity at earlier timepoints likely reflects differences in the initial stroke compositions, with *b* beginning with a downward vertical stroke and *s* beginning with a counterclockwise arc. Finally, *S*_*o,x*_ exhibits a complete absence of similarity, indicating no shared neural dynamics, consistent with the absence of shared stroke components in the characters *o* and *x*.

These results are evidence for a compositional organization of neural activity during handwriting. The presence of similar neural activity at different temporal locations across letters indicates that these motifs are not tied to specific character identities. Instead, neural activity appears to be flexibly employed in accordance with the underlying stroke composition of each character.

### B. Improved Generalization with Compositional Decoding

Motivated by the compositional structure observed in neural activity, we evaluated whether explicitly incorporating this structure in neural decoders improves model performance.

While the primary focus of this study is compositionality in handwriting, we also evaluated MOTIF on attempted speech data. This allowed us to assess the generality of our proposed compositional decoding framework across behavioral modalities (speech and handwriting) and cortical regions (hand-knob area and premotor cortex).

We compared MOTIF with a baseline model that predicts the behavior class without any motif-level supervision. The baseline model follows the same overall architecture as the compositional model but excludes the motif prediction branch. Performance evaluations are conducted between MOTIF and baseline models with matched dimensionalities in the TCN backbone and the behavior prediction branch. We separately train and evaluate the MOTIFmodel on handwriting and speech data under two data regimes: a 50-50 test-train split and a 2-shot generalization test.

**TABLE 1.**
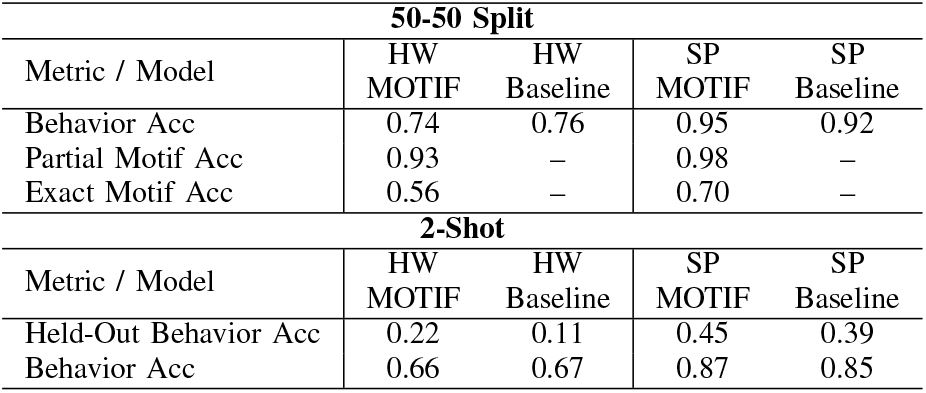
Performance comparison between MOTIF and Baseline models, evaluated on Handwriting (HW) and Speech (SP) datasets. The reported accuracies are on the test split. HO stands for held-out class.

Under the 50-50 test-train split, both models achieved comparable behavior-decoding accuracy (Tab. I). Besides matching the baseline model’s behavior-prediction accuracy, MOTIF also recovered compositional motifs (strokes, phonemes) for the behaviors (characters, words).

In the 2-shot generalization test, MOTIF consistently outperforms the baseline model (Tab. I). Improvements in performance were more pronounced in the held-out class, highlighting MOTIF’s generalization capabilities. Motif-level evaluation corroborated this trend, with a Jaccard score of 0.62 between the model’s predictions and ground truth, indicating that, on average, nearly 2 out of 3 strokes of the held-out character were correctly predicted.

This is significant because compositional decoding in MOTIF enables reliable decoding of sparsely observed behaviors, thereby addressing the imbalance inherent in BCI data, where many behavioral classes have few samples [22], [25], [26].

### C. Interpretability of MOTIF via Attention Analysis

To gain interpretability into the compositional model, we analyzed the attention weights learned by the motif prediction branch of MOTIF. Visualization of the attention weights highlights temporal segments of neural activity that contribute strongly to the prediction of a given motif (Fig. 4).

**Fig. 4.**
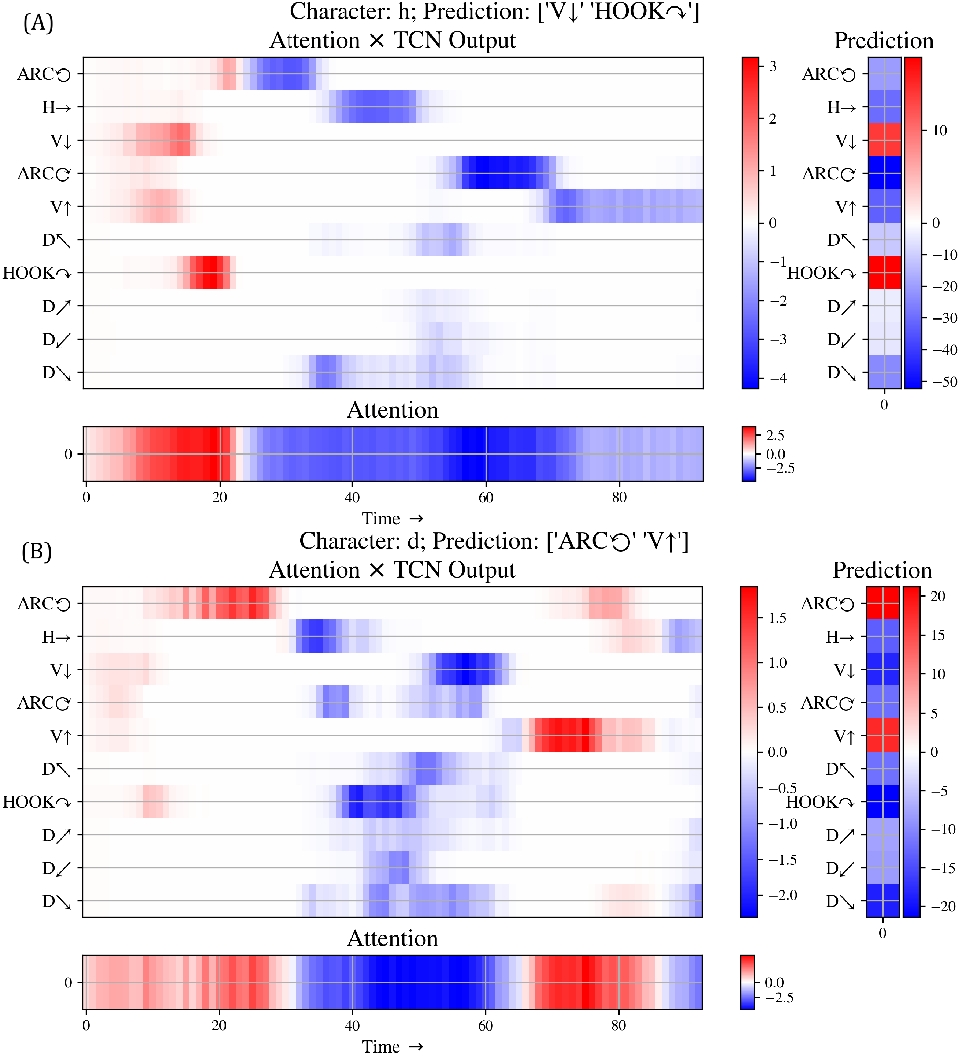
Attention Analysis. Visualization of the motif prediction (right), attention weights (bottom), and attention pooled (center) features obtained from MOTIF for letter ‘h’ and ‘d’. Only positive values (red) contribute towards the motif prediction.

These attention weights exhibit temporal localization within individual trials and correspond to identifiable stroke sequences. The temporal ordering of the attention peaks reflected the ordering of stroke components in the character. For instance, the attention pooled features obtained from MOTIF for the letter ‘h’ (Fig. 4, A) predict the vertical-down (↓) and hook strokes (↷), in that order. Similarly for the letter ‘d’, the attention pooled features first predict the anti-clockwise (⟲) stroke, followed by the vertical-up (↑) stroke. This structure emerged despite the model being trained only to learn the compositional set of motifs without explicit supervision on the stroke sequence. The recovery of sequential structure highlights MOTIF’s ability to leverage compositional neural dynamics, while enhancing interpretability and generalizability.

## IV. Conclusions and Discussion

In this work, we show that the neural activity underlying attempted handwriting exhibits compositional structure and that explicitly modeling this structure in neural decoders enhances generalization and sample efficiency. Our neural similarity analysis revealed distinct bands of elevated similarity among characters that share common stroke components. Importantly, the temporal localization of these bands varied across letter pairs, reflecting differences in stroke ordering and duration of the strokes. Our results indicate that the neural representations of strokes are independent of character identities and are instead flexibly leveraged to compose characters.

We proposed MOTIF, a temporal decoding framework that leverages compositionality through supervision at the motif- and behavioral-levels. Under a standard 50-50 test-train split, the performance of MOTIF and baseline models was comparable, with MOTIF additionally learning the compositional structure in the behavior. In the two-shot generalization test MOTIF far outperformed the baseline model on the heldout class (6% for speech, 11% for handwriting). These results demonstrate that compositional supervision provides an effective inductive bias for BCI decoders. By encouraging the reuse of motif-level neural representations across behaviors, MOTIF achieved substantial improvement in generalization in low-data regimes. The proposed approach offers a promising solution to a key challenge for BCI decoders posed by highly non-uniform behavioral distributions, in which many behaviors are underrepresented by limited samples [22], [25], [26].

Furthermore, analysis of the learned MOTIF attention weights provided an interpretable view into how temporal neural information is leveraged for decoding. Across characters, the temporal localization of attention weights reflected the stroke order of each character. Notably, this structure was discovered implicitly as the model was trained only to identify the set of compositional strokes, without any supervision on the stroke sequences. This approach complements neural latent–based methods [8] by revealing potential transitions in behavioral states and additionally inferring related motif labels. Having this interpretable view of a BCI decoder is particularly valuable as it provides a principled approach for model validation and user-specific adaptations.

In addition to analyzing compositional structure in handwriting neural activity, we also evaluated MOTIF on neural activity underlying attempted speech [2]. This allowed us to test MOTIF on neural activity recorded from a different cortical region, and consider a behavior that is widely regarded to be compositional at the phoneme-level. Our results show that the proposed compositional decoding framework generalizes across behavior modalities (handwriting and speech) and brain regions (hand-motor cortex and premotor cortex).

As BCIs move towards real-world deployment, the ability to decode novel behaviors given limited prior information about the overall behavior becomes essential. By leveraging the compositional structure in neural activity, MOTIF provides a principled step towards decoding behaviors in naturalistic settings.

## Notes

This work was supported by NSF award EFRI 2223822.

### Competing Interest Statement

VG holds shares in Neuralink Corp. and is Chief Scientific Officer and an options holder at Paradromics, Inc.

## References

[1] F. R. Willett, D. T. Avansino, L. R. Hochberg, J. M. Henderson, and K. V. Shenoy, “High-performance brain-to-text communication via handwriting,” Nature, vol. 593, no. 7858, pp. 249–254, 2021.

[2] F. R. Willett, E. M. Kunz, C. Fan, D. T. Avansino, G. H. Wilson, E. Y. Choi, F. Kamdar, M. F. Glasser, L. R. Hochberg, S. Druckmann, K. V. Shenoy, and J. M. Henderson, “A high-performance speech neuroprosthesis,” Nature, vol. 620, no. 7976, pp. 1031–1036, 2023.

[3] S. L. Metzger, K. T. Littlejohn, A. B. Silva, D. A. Moses, M. P. Seaton, R. Wang, M. E. Dougherty, J. R. Liu, P. Wu, M. A. Berger, I. Zhuravleva, A. Tu-Chan, K. Ganguly, G. K. Anumanchipalli, and E. F. Chang, “A high-performance neuroprosthesis for speech decoding and avatar control,” Nature, vol. 620, no. 7976, pp. 1037–1046, 2023.

[4] N. S. Card, M. Wairagkar, C. Iacobacci, X. Hou, T. Singer-Clark, F. R. Willett, E. M. Kunz, C. Fan, M. Vahdati Nia, D. R. Deo, A. Srinivasan, E. Y. Choi, M. F. Glasser, L. R. Hochberg, J. M. Henderson, K. Shahlaie, S. D. Stavisky, and D. M. Brandman, “An accurate and rapidly calibrating speech neuroprosthesis,” New England Journal of Medicine, vol. 391, no. 7, pp. 609–618, 2024.

[5] M. Wairagkar, N. S. Card, T. Singer-Clark, X. Hou, C. Iacobacci, L. M. Miller, L. R. Hochberg, D. M. Brandman, and S. D. Stavisky, “An instantaneous voice-synthesis neuroprosthesis,” Nature, pp. 1–8, 2025.

[6] N. Bernstein, The Co-ordination and Regulation of Movements. Pergamon Press, 1967. [Online]. Available: https://books.google.com/books?id=mUhzjwEACAAJ

[7] S. F. Giszter, “Motor primitives—new data and future questions,” Current opinion in neurobiology, vol. 33, pp. 156–165, 2015.

[8] A. J. Zimnik and M. M. Churchland, “Independent generation of sequence elements by motor cortex,” Nature neuroscience, vol. 24, no. 3, pp. 412–424, 2021.

[9] T. Wang, Y. Chen, Y. Zhang, and H. Cui, “Multiplicative joint coding in preparatory activity for reaching sequence in macaque motor cortex,” Nature Communications, vol. 15, no. 1, p. 3153, 2024.

[10] N. P. Shah, D. Avansino, F. Kamdar, C. Nicolas, A. Kapitonava, C. Vargas-Irwin, L. R. Hochberg, C. Pandarinath, K. V. Shenoy, F. R. Willett, and J. M. Henderson, “Pseudo-linear summation explains neural geometry of multi-finger movements in human premotor cortex,” Nature Communications, vol. 16, no. 1, p. 5008, 2025.

[11] A. Maes, M. Barahona, and C. Clopath, “Learning compositional sequences with multiple time scales through a hierarchical network of spiking neurons,” PLoS computational biology, vol. 17, no. 3, p. e1008866, 2021.

[12] M. S. Willsey, N. P. Shah, D. T. Avansino, N. V. Hahn, R. M. Jamiolkowski, F. B. Kamdar, L. R. Hochberg, F. R. Willett, and J. M. Henderson, “A high-performance brain–computer interface for finger decoding and quadcopter game control in an individual with paralysis,” Nature Medicine, vol. 31, no. 1, pp. 96–104, 2025.

[13] D. A. Moses, S. L. Metzger, J. R. Liu, G. K. Anumanchipalli, J. G. Makin, P. F. Sun, J. Chartier, M. E. Dougherty, P. M. Liu, G. M. Abrams, A. Tu-chan, K. Ganguly, and E. F. Chang, “Neuroprosthesis for decoding speech in a paralyzed person with anarthria,” New England Journal of Medicine, vol. 385, no. 3, pp. 217–227, 2021.

[14] The CMU pronouncing dictionary, “CMUdict.” [Online]. Available: http://www.speech.cs.cmu.edu/cgi-bin/cmudict

[15] C. Lea, R. Vidal, A. Reiter, and G. D. Hager, “Temporal convolutional networks: A unified approach to action segmentation,” in European conference on computer vision. Springer, 2016, pp. 47–54.

[16] S. Bai, Z. Kolter, and V. Koltun, “An empirical evaluation of generic convolutional and recurrent networks for sequence modeling,” arXiv preprint 1803.01271, 2018.

[17] H. Temmar, M. S. Willsey, J. T. Costello, M. J. Mender, L. H. Cubillos, J. C. DeMatteo, J. L. Lam, D. M. Wallace, M. M. Kelberman, P. G. Patil, and C. A. Chestek, “Investigating the benefits of artificial neural networks over linear approaches to BMI decoding,” Journal of Neural Engineering, 2025.

[18] J. Huang, P. Tostado-Marcos, S. M. Narasimha, A. N. Patel, E. M. Arneodo, T. Q. Gentner, G. Mishne, and V. Gilja, “Guiding brain-to-vocalization decoder design using structured generalization error,” in 2024 46th Annual International Conference of the IEEE Engineering in Medicine and Biology Society (EMBC). IEEE, 2024, pp. 1–4.

[19] V. Feldman, “Does Learning Require Memorization? A Short Tale about a Long Tail,” in Proceedings of the 52nd Annual ACM SIGACT Symposium on Theory of Computing, 2020, pp. 954–959.

[20] V. Feldman and C. Zhang, “What Neural Networks Memorize and Why: Discovering the Long Tail via Influence Estimation,” Advances in Neural Information Processing Systems, vol. 33, pp. 2881–2891, 2020.

[21] Z. Liu, Z. Miao, X. Zhan, J. Wang, B. Gong, and X. Y. Stella, “Open Long-Tailed Recognition in a Dynamic World,” IEEE Transactions on Pattern Analysis and Machine Intelligence, vol. 46, no. 3, pp. 1836– 1851, 2022.

[22] J. Huang, A. N. Patel, S. M. Narasimha, G. Mishne, and V. Gilja, “Word-level error analysis in decoding systems: From speech recognition to brain-computer interfaces,” in Proc. Interspeech 2025, 2025, pp. 5563– 5567.

[23] A. Paszke, S. Gross, F. Massa, A. Lerer, J. Bradbury, G. Chanan, T. Killeen, Z. Lin, N. Gimelshein, L. Antiga, A. Desmaison, A. Kopf, E. Yang, Z. DeVito, M. Raison, A. Tejani, S. Chilamkurthy, B. Steiner, L. Fang, J. Bai, and S. Chintala, “Pytorch: An imperative style, high-performance deep learning library,” Advances in neural information processing systems, vol. 32, 2019.

[24] D. P. Kingma and J. Ba, “Adam: A method for stochastic optimization,” arXiv preprint 1412.6980, 2014.

[25] P. Thülke, Y.-J. Mantilla-Ramos, H. Abdelhedi, C. Maschke, A. Dehgan, Y. Harel, A. Kemtur, L. M. Berrada, M. Sahraoui, T. Young, A. B. Pepin, C. E. Khantour, M. Landry, A. Pascarella, V. Hadid, E. Combrisson, J. O’Byrne, and K. Jerbi, “Class imbalance should not throw you off balance: Choosing the right classifiers and performance metrics for brain decoding with imbalanced data,” NeuroImage, vol. 277, p. 120253, 2023.

[26] T. Liu and D. Yang, “A three-branch 3d convolutional neural network for eeg-based different hand movement stages classification,” Scientific Reports, vol. 11, no. 1, p. 10758, 2021.

